# Thermal acclimation of leaf respiration consistent with optimal plant function

**DOI:** 10.1101/434084

**Authors:** Han Wang, Owen K. Atkin, Trevor F. Keenan, Nicholas Smith, Ian J. Wright, Keith J. Bloomfield, Jens Kattge, Peter B. Reich, I. Colin Prentice

**Affiliations:** Ministry of Education Key Laboratory for Earth System Modelling, Department of Earth System Science, Tsinghua University, Beijing 100084, China; Joint Centre for Global Change Studies, Beijing 100875, China; Division of Plant Sciences, Research School of Biology, The Australian National University, Canberra, ACT 2601, Australia; Australian Research Council Centre of Excellence in Plant Energy Biology, Research School of Biology, The Australian National University, Canberra, ACT 2601, Australia; Department of Environmental Science, Policy and Management, UC Berkeley, Berkeley, CA, USA; Climate and Ecosystem Sciences Division, Lawrence Berkeley National Laboratory, Berkeley, CA, USA; Department of Biological Sciences, Texas Tech University, Lubbock, TX, USA; Department of Biological Sciences, Macquarie University, NSW 2109, Australia; Max Planck Institute for Biogeochemistry, Jena, Germany; German Center for Integrative Biodiversity Research Halle-Jena-Leipzig, Leipzig, Germany; Department of Forest Resources, University of Minnesota, St. Paul, MN 55108, USA; Hawkesbury Institute for the Environment, Western Sydney University, Penrith, NSW 2753, Australia; AXA Chair of Biosphere and Climate Impacts, Department of Life Sciences, Imperial College London, Silwood Park Campus, Buckhurst Road, Ascot SL5 7PY, UK

## Main

### Introduction

Terrestrial plant respiration is a major component of the global carbon cycle, releasing *ca.* 60 Pg C yr^−1^ to the atmosphere: six times more than anthropogenic CO_2_ emissions from all sources combined (Ciais *et al.* 2014). About half this flux is leaf maintenance respiration in darkness (*R*_d_) (Atkin *et al.* 2007). *R*_d_ is highly temperature-dependent, following a near-exponential relationship over short time-scales (Heskel *et al.* 2016). It has been predicted that global warming will increase *R*_d_ and accelerate future climate change via a carbon-climate feedback (Cox *et al.* 2000; Huntingford *et al.* 2013). However, large uncertainties in future carbon cycle responses to warming persist, likely because model formulations of the temperature responses of photosynthesis and *R*_d_ remain poorly constrained by both theory and observations (Ziehn *et al.* 2011; Booth *et al.* 2012; Friedlingstein *et al.* 2014).

Representations of *R*_d_ in most land surface models (LSMs) are based on the instantaneous temperature response of *R*_d_, governed by enzyme kinetics, relative to a baseline rate, typically at 25°C (*R*_d,25_) (Atkin *et al.* 2017). *R*_d,25_ is commonly assumed to be proportional either to area-based leaf nitrogen content (*N*_area_), or to the maximum rate of carboxylation at 25°C (*V*_cmax,25_) (Rogers 2014). Predicting photosynthetic traits from *N*_area_ is problematic because a large fraction of leaf N is contained in cell walls (Onoda *et al.* 2017), and significant fractions are also allocated to other non-photosynthetic functions including defense, storage and osmoregulation (Dong *et al.*, 2017). Many models treat *V*_cmax,25_ and *R*_d,25_ as constant for each plant functional type (PFT), but spatial and temporal trait variations within PFTs are to be expected (Wang *et al.* 2017b) and have been reported (Kattge *et al.* 2011, Atkin et al. 2015). Moreover, trait differences observed among PFTs could be caused by acclimation to different environments, rather than intrinsic properties of PFTs.

Experiments have shown acclimation of *R*_d_, such that its response to growth temperature over a week or longer is shallower than its response to temperature variation in the short term (Atkin & Tjoelker 2003; Gifford 2003; Smith & Dukes 2013; Aspinwall *et al.* 2016; Drake *et al.* 2016; Reich *et al.* 2016; Scafaro *et al.* 2017). The acclimation of *R*_d_ to growth temperature is also evident in spatial observations (Atkin *et al.* 2015; Slot & Kitajima 2015), showing a far weaker pole-to-equator gradient than would be expected if field-measured *R_d_* followed the instantaneous response of *R*_d_ to temperature. By analysing the temperature responses of *R*_d_ in different datasets, Vanderwel *et al.* (2015) demonstrated consistency between the observed spatial pattern of *R*_d_ and the acclimation of *R*_d_ over time. Similar levels of thermal acclimation of photosynthesis and respiration have been shown to occur in different PFTs (Campbell *et al.* 2007; Smith & Dukes 2017b). Pervasive *R*_d_ acclimation implies a weaker positive carbon-climate feedback than implied by the temperature response of enzyme kinetics (i.e. the instantaneous response) (Reich *et al.* 2016; Smith *et al.* 2016; Huntingford *et al.* 2017). Neglecting acclimation in LSMs is thus a potential major source of bias in Earth system model predictions (<u>Smith & Dukes, 2013</u>), as recently demonstrated by Huntingford *et al.* (2017).

Still missing is a theoretical explanation for *R*_d_ acclimation. Conclusions from empirical studies alone (Wright *et al.* 2006) remain subject to the limitations of sampled region, observational period and experimental design. A theoretical basis is essential to build confidence in carbon-cycle predictions (Prentice *et al.*, 2015). As advocated e.g. by Marquet *et al.* (2014), theories grounded in first principles have the potential to generate explicit quantitative predictions with few assumptions or unconstrained parameters, providing independent standards for comparison with empirical data. Here, we develop such a theory for *R*_d_ acclimation, based on the assumption that the various metabolic functions of *R*_d_ are coordinated with photosynthetic capacity, indexed by *V*_cmax_. In combination with predictions of optimally acclimated *V*_cmax_ based on the coordination hypothesis (Chen et al., 1993; Haxeltine & Prentice, 1996; Maire et al., 2012; Togashi *et al.*, 2018) – whereby optimal *V*_cmax_ is just sufficient to use available resources under current average environmental conditions – our approach allows us to formulate and test the sensitivities of the traits of interest (*R*_d_, *V*_cmax_) to temperature, yielding results applicable to all plants.

Our theory is based on the concept of eco-evolutionary optimality, which derives from the premise that natural selection favours efficient resource allocation by eliminating unsuccessful or uncompetitive trait combinations (Givnish 1986a; Tilman 1999). Optimality-based theories have proven predictive power for plant functions including leaf venation networks (Blonder *et al.* 2017), stomatal behaviour (Cowan 1986; Givnish 1986b; Farquhar *et al.* 2002; Lin *et al.* 2015; Wolf *et al.* 2016; Dewar *et al.* 2018), leaf-level CO_2_ drawdown (Prentice *et al.* 2014), phenology (Kikuzawa *et al.* 2013; Xu *et al.* 2017), leaf nitrogen content (Wright *et al.* 2003; Maire *et al.* 2012; Dong *et al.* 2017) and adaptations to elevation (Wang *et al.* 2017a); and ecosystem processes including vegetation succession (Weng *et al.* 2017) and primary production (Keenan *et al.* 2016; Wang *et al.* 2017b). By linking *R*_d_ with optimal *V*_cmax_ acclimation (Box 1), we derive quantitative predictions of the thermal sensitivities of acclimated *R*_d_ and *V*_cmax_ evaluated at the prevailing growth temperature, and at 25°C. Our theory implies that correlations between *V*_cmax,25_ and *N*_area_, and between *R*_d,25_ and *N*_area_ primarily reflect the N requirements of metabolism (as implied by the coordination hypothesis) rather than ‘N limitation’ of either *V*_cmax_ or *R*_d_ – that is, the amount of ‘metabolic’ N in the leaf is optimized for current conditions. These predictions are tested using two extensive field observational datasets (Atkin *et al.*, 2015; Smith & Dukes, 2017).

### Materials and Methods

#### Theoretical framework

Leaf dark respiration (*R*_d_) is closely coupled with photosynthetic activity (Hoefnagel *et al.* 1998; Wright *et al.* 2004; Noguchi & Yoshida 2008; Tcherkez 2012). As described by the standard biochemical model of photosynthesis (Farquhar *et al.* 1980), the instantaneous rate of photosynthesis by C_3_ plants is limited either by the capacity of the enzyme Ribulose-1,5-bisphosphate (RuBP) carboxylase/oxygenase (Rubisco) for the carboxylation of RuBP (*V*_cmax_), or by the rate of electron transport for the regeneration of RuBP, which depends on absorbed light and the electron transport capacity (*J*_max_). *R*_d_ is used to support diverse metabolic processes including protein turnover, phloem loading, the maintenance of ion gradients between cellular compartments, nitrate reduction, and the turnover of phospholipid membranes. Among these, protein turnover is the largest contributor to *R*_d_ variation and is expected to scale closely with *V*_cmax_, which sets the daily maximum photosynthetic rate achieved by leaves under natural growing conditions (Amthor 2000; Atkin *et al.* 2000; Cannell & Thornley 2000; Bouma 2005).

We start from the assumption that at the prevailing growth temperature, *V*_cmax_ is proportional to the overall metabolic activity of the leaf, which is supported by *R*_d_. The dimensionless factor *b* (main text equation (1)) is expected to be constant (for all plants) at a given growth temperature. The value of *b* at 25°C (*b*_25_) has been given as 0.011 (Farquhar *et al.*, 1980) or 0.015 (Collatz *et al.* 1991). A constant *b*_25_ implies that a certain quantity of respiratory enzymes is required for the maintenance of a certain quantity of Rubisco. Our initial attention therefore focuses on the prediction of optimal *V*_cmax_ achieved by the acclimation of photosynthetic processes.

We hypothesize that *V*_cmax_ of leaves at any canopy level acclimates to the current environment so as to be just sufficient to allow exploitation of the average available light. This is the “strong form” (Togashi *et al.*, 2018) of the coordination hypothesis (Chen et al., 1993; Haxeltine & Prentice, 1996; Maire et al., 2012) (contrasted with the “weak form” that assumes that the total metabolic N content of the leaf is prescribed, so only the allocation of N to carboxylation versus electron transport capacities is optimized). The hypothesis leads to a prediction of the ecophysiologically relevant (acclimated) values of both *R*_d_ and *V*_cmax_ at the prevailing temperature, i.e. the average growth temperature (*T*_g_). Values of *R*_d_ and *V*_cmax_ at 25°C (*R*_d,25_ and *V*_cmax,25_) are related to quantities of enzymes, while values at growth temperature (*R*_d,tg_ and *V*_cmax,tg_) are hypothesized to be optimized by the plant. Numerical conversion between values applying to different temperatures is accomplished by applying functions (Kattge & Knorr, 2007; Heskel *et al.*, 2016) that describe how the instantaneous rates increase with temperature due to enzyme kinetics. To achieve the same level of *V*_cmax_, a smaller quantity of Rubisco (and therefore a smaller *V*_cmax,25_) is required at a higher temperature (Fig. 1). The coordination hypothesis predicts acclimated values of *R*_d,tg_ and *V*_cmax,tg_ that increase with growth temperature, but less steeply than the responses of enzyme kinetics (Togashi *et al*., 2018). As a result, acclimated values of *R*_d,25_ and *V*_cmax,25_ are predicted to *decline* with growth temperature. Moreover, because the instantaneous responses of *R*_d_ and *V*_cmax_ to temperature are slightly different, these differences have to be compensated by differences in the acclimated responses of *R*_d,tg_ and *V*_cmax,tg_.

**Figure 1:**
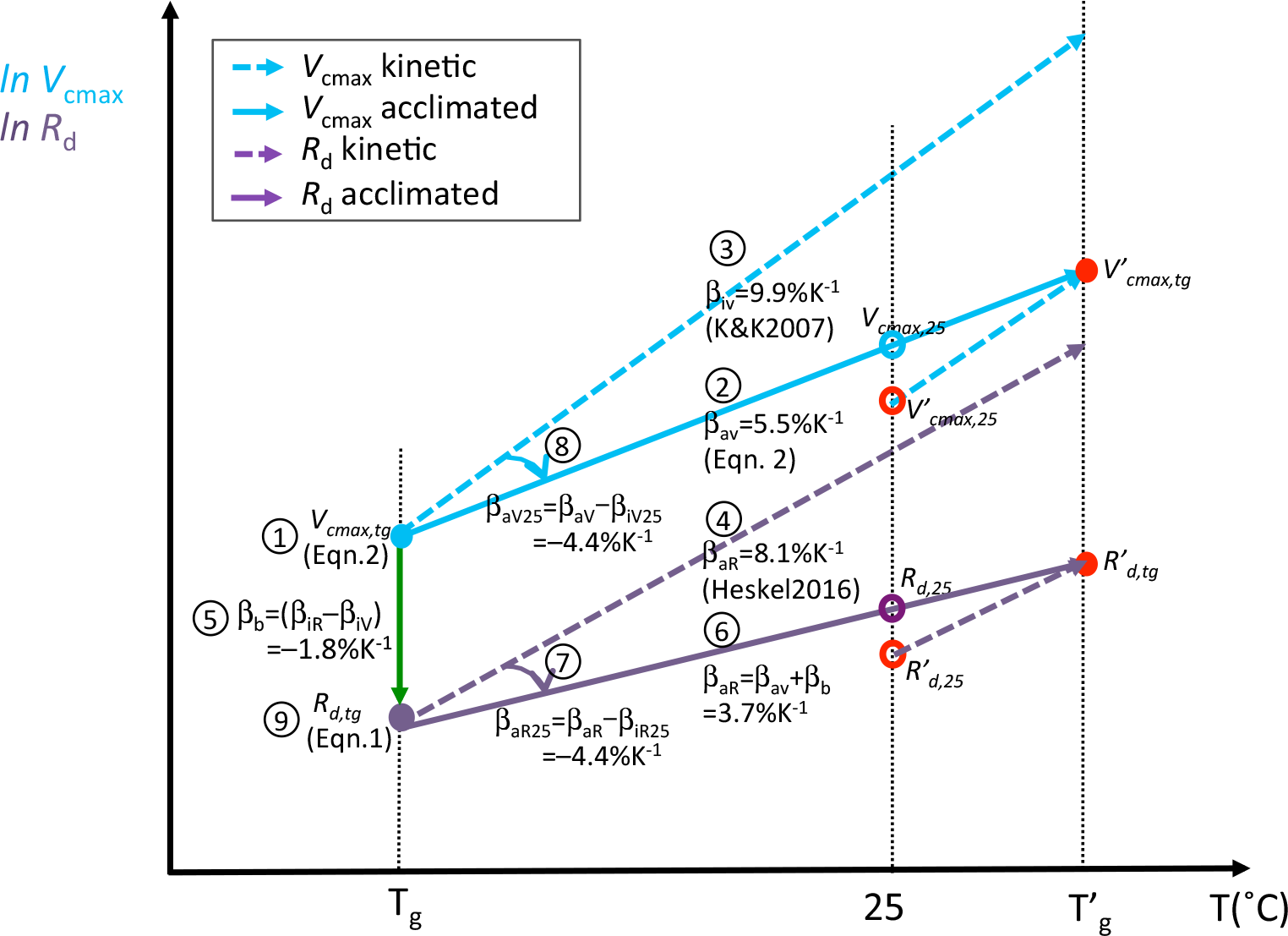
Schematic of the thermal sensitivities of area-based maximum capacity of carboxylation (*V*_cmax_) and leaf dark respiration (*R*_d_). *β*_aV_: the acclimated thermalsensitivity of *V*_cmax_ assessed at growth temperature (*T*_g_). *β*_aR_: the acclimated thermal sensitivity of *R*_d_ assessed at growth temperature. *β*_iV_: the kinetic thermal sensitivity of *V*_cmax_ assessed at growth temperature. *β*_iR_: the kinetic thermal sensitivity of *R*_d_ assessed at growth temperature. *β*_aV25_: the acclimated thermal sensitivity of *V*_cmax_ assessed at 25°C. *β*_aR25_: the acclimated thermal sensitivity of *R*_d_ assessed at 25°C. *β*_b_: the thermal sensitivity of the ratio of *R*_d_ to *V*_cmax_ assessed at growth temperature. *V*_cmax_ and *R*_d_ are assessed at growth temperature (*V*_cmax,tg_ and *R*_d,tg_) as a starting point, 25°C as the common practice (*V*_cmax,25_ and *R*_d,25_) and a higher growth temperature (*T*_g_’) for the acclimation behaviour (*V*’_cmax,tg_, *R*’_d,tg_, *V*’_cmax,25_ and *R*’_d,25_) to warming.

The reasoning set out above can be represented mathematically, leading to a theoretical derivation of the thermal sensitivities of *R*_d_, *V*_cmax_ and their ratio (assessed at both *T*_g_ and 25°C) as follows. We first use the coordination hypothesis to predict the fractional sensitivity of *V*_cmax,tg_ to growth temperature (denoted *β*_aV_, the subscripts *a* and *i* here represent the acclimated and instantaneous responses, respectively), which is predicted to be less than the instantaneous response, i.e. *β*_aV_ < *β*_iV_. The hypothesized proportionality between *R*_d_ and *V*_cmax_ then links the thermal sensitivity of *R*_d,tg_ (*β*_aR_) with *V*_cmax,tg_, since *β*_aR_ = *β*_b_ + *β*_aV_. Here *β*_b_ quantifies the variation of *b* with temperature, which is due to the slight lower instantaneous thermal responses of *R*_d_ (*β*_iR_) than *V*_cmax_, i.e. *β_b_ = β*_iR_ – *β*_iV_ < 0. Making use of the general properties of logarithms, *β*_*a*V25_ and *β*_aR25_ are predicted as a secondary consequence of *V*_cmax,tg_ and *R*_d,tg_ acclimation combined with enzyme kinetics: *β*_aV25_ = *β*_aV_ – *β*_iV_ = *β*_aR25_ = *β*_aR_ − *β*_iR_ < 0

#### Quantitative predictions

Our approach (Box 1) is based on the assumption that acclimated *R*_d_ is proportional to photosynthetic capacity, represented by *V*_cmax_, where their ratio (*b)* may be a function of temperature:

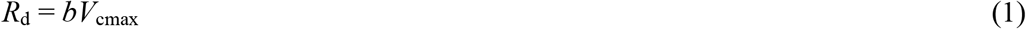

We therefore first explain quantitative predictions of the thermal acclimation of optimal *V*_cmax_.

The coordination hypothesis predicts a spatial and temporal pattern in *V*_cmax_ assessed at growth temperature, reflecting acclimation to the prevailing environmental conditions of an individual leaf. In response to environmental variations, it predicts that *V*_cmax_ can vary vertically within the canopy, geographically among sites, and temporally with atmospheric CO_2_ concentration and climate (Haxeltine & Prentice 1996; Ainsworth & Long 2005; Maire *et al.* 2012). *V*_cmax_ values have been shown experimentally to acclimate to sustained changes in growth temperature, such that *V*_cmax_ assessed at growth temperature (hereafter *V*_cmax,tg_) increases with growth temperature, while *V*_cmax,25_ declines, along with the Rubisco amount and the fraction of leaf N allocated to Rubisco (Scafaro *et al.* 2017).

Quantitative predictions of *V*_cmax_ can be obtained from the co-ordination hypothesis by equating the Rubisco-limited (*A*_C_) and electron transport-limited (*A*_J_) rates of C_3_ photosynthesis in the Farquhar *et al.* (1980) model (Wang *et al.*, 2017b). For simplicity, we shall assume that the light response of *A*_J_ under natural light conditions is effectively linear up to the point at which *A*_C_ becomes limiting, implying that limitation of photosynthesis by *J*_max_ under average field conditions is generally avoided (Wang *et al.* 2014; Keenan *et al.* 2016; Dong *et al.* 2017; Wang *et al.* 2017a; Togashi *et al.* 2018). Thus, under field conditions the coordination hypothesis predicts (Dong *et al.*, 2017; Togashi *et al*., 2018):

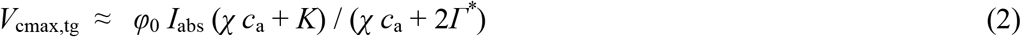

where *φ*_0_ is the intrinsic quantum efficiency of photosynthesis, which is independent of temperature over the normal range of metabolic activity (Collatz *et al.* 1990), and *I*_abs_ is the leaf absorbed photosynthetic photon flux density (PPFD). *χ* is the ratio of leaf-internal to ambient partial pressure of CO_2_, *c_a_* is the ambient partial pressure of CO_2_, *Γ^*^* is the photorespiratory compensation point, and *K* is the effective Michaelis-Menten coefficient of Rubisco. *Γ^*^* and *K* are temperature-dependent following Arrhenius relationships measured e.g. by Bernacchi *et al.* (2001). The least-cost hypothesis (Prentice *et al.*, 2014; Wang *et al.*, 2017b) predicts optimal *χ* as a function of growing-season mean values of temperature (*T*_g_) and vapour pressure deficit (*D*), and elevation (*z*). This prediction is quantitatively supported by worldwide measurements of *χ* across species and environments (Wang *et al.* 2017b). Equation (2) yields estimates of *V*_cmax_ given *χ* and field-relevant average values of *c*_a_, temperature and PPFD. The theoretical temperature dependence of *V*_cmax,tg_ arises from the separate temperature responses of *χ*, *Γ*^*^ and *K*. The sensitivity of *V*_cmax,tg_ to temperature can be obtained by differentiating equation (2). Evaluating the result under standard conditions (*T*_g_ = 25°C, *D* = 1 kPa, *z* = 0, *c*_a_ = 40 Pa) yields *β*_aV_ = 5.5% K^−1^ (*β*_a*V*_ is the fractional sensitivity of *V*_cmax,tg_ to temperature after acclimation: see Box 1 for definitions, Figure 1 for a graphical explanation, and Appendix 1 for derivations). This value derives primarily from the sensitivities of *K* and *Γ*^*\**^ to temperature (8.5% K^−1^ and 5.4% K^−1^, respectively), which depend on their activation energies (Bernacchi *et al.* 2001), and to a lesser extent on the sensitivity of *χ* to temperature (0.9% K^−1^).

*V*_cmax,tg_ is then corrected from *T*_g_ to 25°C using the enzyme-kinetic temperature response of *V_cmax_* (Kattge & Knorr 2007). Evaluated at 20°C (mean *T*_g_ in our dataset), this function yields an instantaneous thermal sensitivity of *V*_cmax_ of *β*_iV_ = 9.9% K^−1^, higher than the acclimated thermal sensitivity (*β*_aV_ = 5.5% K^−1^). The thermal sensitivity of acclimated *V*_cmax,25_ is predicted as the difference between the thermal sensitivities of *V*_cmax_ acclimated to the growth temperature (*V*_cmax,tg_) and the instantaneous enzyme-kinetic response of *V*_cmax_ (Box 1: *β*_aV25_ = *β*_aV_ *– β*_iV_ = −4.4% K^−1^

Heskel *et al.* (2016) provided an estimate of *R*_d_ at a reference temperature (*T*_ref_):

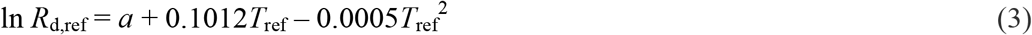

where *a* is an empirical constant varying among biomes, representing the natural log of the value of *R*_d_ at 0°C. The enzyme-kinetic response of *R*_d_ to temperature (*β*_iR_) as given by Heskel *et al.* (2016) is 8.1% K^−1^ at the mean *T*_g_ of the data. The enzyme-kinetic thermal response of *R*_d_ is slightly smaller than *β*_iV_, and leads to a thermal response of parameter *b* in equation (1) given by the difference between *β*_iR_ and *β*_iV_ (*β_b_* = *β*_iR_ − *β*_iV_ = −1.8% K^−1^). This then generates a prediction of *β_aR_* = 3.7% K^−1^ and *β_a_*_R25_ = –4.4%, which is the same as *β*_*a*V25_, consistent with the assumption that *b*_25_ is a constant (Box 1). Derivations are provided in Supporting Information (Appendix 1).

#### Empirical analyses

##### Photosynthesis and respiration data

We combined two *R*_d_ datasets. GlobResp (Atkin *et al.* 2015) contains measurements of *R*_d_, *V*_cmax_, *N*_area_ and leaf mass per area (LMA) from 899 species at 100 locations across the major biomes and continents, including data from an earlier compilation by Wright *et al.* (2004). LCE (Smith & Dukes 2017a) contains field measurements of leaf carbon exchange (including *R*_d_ and *V_cmax_*) and leaf chemical traits (including *N*_area_ and LMA) from 98 species at 12 locations spanning 53° latitude in North and Central America (Fig. S1). *R*_d_ measurements in both datasets followed the same protocol. Both were taken on fully expanded leaves in daytime after a period of dark adjustment. Inhibition of respiration in the light was not assessed. *V*_cmax_ values in GlobResp were estimated by the ‘one-point method’ whereas those in LCE were estimated from full *A*-*c*_i_ curves; these methods give closely similar results (De Kauwe *et al.* 2016). Replicated measurements in LCE on the same species and site were averaged. Juvenile samples were excluded. We index *T*_g_ by the mean temperature during the thermal growing season when temperatures are above 0°C, mGDD_0_ (Harrison *et al.* (2010). *V*_cmax_ and *R*_d_ values in both datasets are provided with information about measurement temperatures. The values were adjusted both to mGDD_0_ and to 25°C using the relevant kinetic responses, as given by (Kattge & Knorr 2007) and Heskel *et al.* (2016). A global climatology of monthly temperature provided by the Climatic Research Unit at a grid resolution of 10 arc minutes (CRU CL2.0) (New *et al.* 2000) was used to provide estimates of mGDD_0_ for each location. Thermal acclimation of *R*_d_ should apply to both C_3_ and C_4_ plants, but our theoretical prediction of *V*_cmax_ acclimation here is developed for C_3_ plants, and we did not include C_4_ species in our analysis.

##### Statistical analysis

To test our predictions of the acclimated thermal sensitivities of *R*_d_ and *V*_cmax_ quantitatively, the *R*_d_ and *V*_cmax_ data (assessed at mGDD_0_ and 25°C) were first normalized with site-mean PPFD absorbed by leaves (PPFD_L_) before performing the Ordinary Least Squares (OLS) regression against temperature. This normalization is appropriate because *V*_cmax_ is both predicted (see Appendix 1) and observed to vary in proportion to PPFD (Niinemets & Keenan, 2012). If it were omitted, the positive effect of PPFD on *R*_d_ and *V*_cmax_ would contribute to the fitted slope of mGDD_0_ due to the strong correlation between those two variables (Fig. S2). Site-mean PPFD_L_ was estimated from growing-season total incident PPFD at the top of the canopy (PPFD_0_):

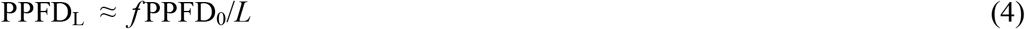

where *f* is the fraction of incident PPFD absorbed by the canopy (from SeaWiFS data: (Gobron *et al.* 2006; Kelley *et al.* 2013) and *L* is the leaf area index estimated from Beer’s law:

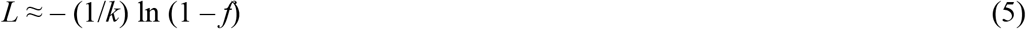

with *k* = 0.5 (Dong *et al.* (2017). This general approximation is used because we do not have information on the light levels of species occupying different canopy strata. PPFD_0_ was calculated from CRU CL2.0 data using SPLASH (Davis *et al.* 2017). We applied OLS linear regression of normalized (and natural log-transformed) *R*_d_ and *V*_cmax_ values against mGDD_0_. The resulting slope coefficients are directly comparable with the thermal sensitivities predicted by theory. To check the impact of the PPFD normalization, we also performed regressions without it. Thermal acclimation of *R*_d_ and *V*_cmax_ was further tested within PFTs, with species assigned to deciduous and evergreen needleleaf and broadleaf trees, deciduous and evergreen shrubs, and C3 herbaceous plants.

To test the hypothesis that *R*_d_ is mainly determined by *V*_cmax_ rather than leaf or soil characteristics, we also included LMA and soil pH as additional predictors in the OLS regression above. LMA carries information on the structural component of plant leaves. Broadly speaking, higher soil pH indicates higher soil fertility (Jenny, 1994; Sinsabaugh & Follstad Shah, 2012), and pH has been shown to influence *χ* (Wang et al. 2017b). These two covariates were selected to test any potential influences of leaf structure and soil nutrient availability, respectively, on *R*_d_. An estimate of soil pH for each location was extracted from the Harmonized World Soil Database (http://www.iiasa.ac.at). We applied Ordinary Least Squares (OLS) linear regression of *R*_d_ versus *V*_cmax_ (standardized to 25°C and to mGDD_0_, without transformation) to estimate *b*_25_ and *b* directly from the fitted slopes. Relationships of *N*_area_ with *V*_cmax_, *R*_d_, and LMA were also tested by OLS multiple linear regression.

### Results

#### Sensitivity of acclimated R_d_ and V_cmax_ to growth temperature

The predicted sensitivity of *V*_cmax_ to growth temperature (*β*_aV_) was 5.5% K^−1^ under standard environmental conditions. This value is identical with the fitted regression coefficient of normalized and transformed *V*_cmax,tg_ against mGDD_0_ (5.6% ± 0.3%: mean and 95% confidence interval, *R*^2^ = 0.50) (Table 1, Fig. 2).

**Table 1:** Summary of Ordinary Least-Squares regressions for natural log-transformed area-based leaf dark respiration (*R*_d_), area-based maximum carboxylation rate (*V*_cmax_) and their ratio as a function of growth temperature. Both *R*_d_ and *V*_cmax_ have been standardized to growth temperature (*R*_d,tg_ and *V*_cmax,tg_) and to 25 °C (*R*_d,25_ and *V*_cmax,25_), and normalized by site-mean leaf absorbed photosynthetic photon flux density. The fitted coefficient and its confidence interval are shown together with the corresponding theoretical expectation, the intercept (mean ± standard error), the coefficient of determination (*r^2^*) and the degrees of freedom (*df*). Non-significant coefficients are shown in grey.

| Quantity | Theoretical value | Fitted coefficient | Confidence intervals | | Intercept | $r^2$ | $df$ |
| --- | --- | --- | --- | --- | --- | --- | --- |
|  |  |  | 2.5% | 97.5% |  |  |  |
| $R_{d,\text{tg}}$ | 0.037 | 0.033 | 0.029 | 0.038 | -9.335±0.051 | 0.34 | 1245 |
| $V_{\text{cmax,tg}}$ | 0.055 | 0.056 | 0.050 | 0.061 | -6.386±0.054 | 0.50 | 1009 |
| $R_{d,\text{tg}}/V_{\text{cmax,tg}}$ | -0.018 | -0.020 | -0.026 | -0.015 | -2.916±0.060 | 0.05 | 1007 |
| $R_{d,25}$ | -0.044 | -0.049 | -0.054 | -0.045 | -7.261±0.052 | 0.25 | 1245 |
| $V_{\text{cmax},25}$ | -0.044 | -0.044 | -0.049 | -0.038 | -3.907±0.054 | 0.26 | 1009 |
| $R_{d,25}/V_{\text{cmax},25}$ | 0 | -0.005 | -0.010 | 0.001 | -3.317±0.060 | 0.00 | 1007 |

**Figure 2:**
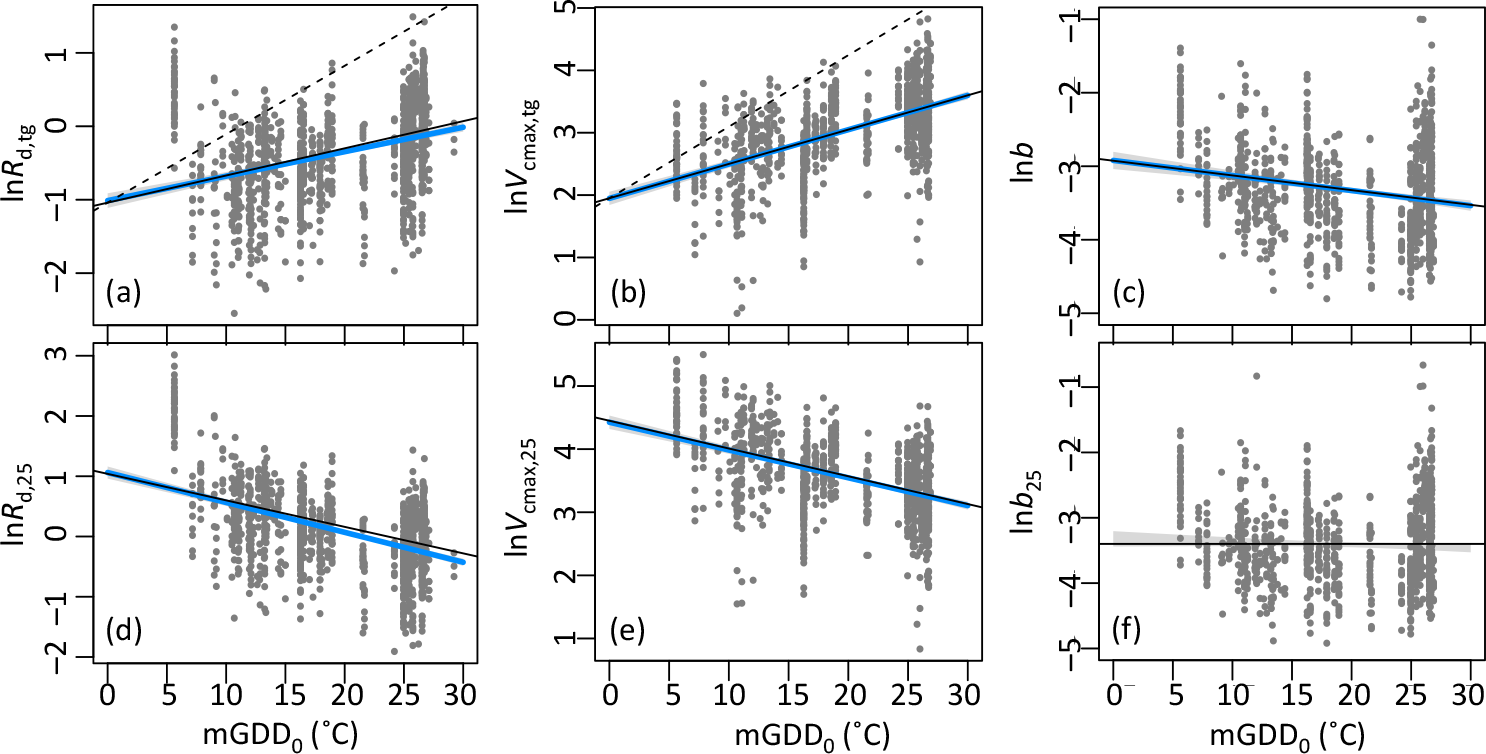
Natural log-transformed light-normalized leaf dark respiration (*R*_d_), maximum carboxylation rates (*V*_cmax_) and their ratio, standardized to growth temperature (*R*_d,tg_, *V*_cmax,tg_, *b*: *R*_d,tg_/*V*_cmax,tg_) and to 25°C (*R*_d,25_, *V*_cmax,25_, *b*_2_: *R*_d,25_/*V*_cmax,25_), as a function of growth temperature (mGDD_0_). Solid blue lines are the fitted lines from Ordinary Least Squares regressions. Solid black lines are theoretical predictions. Dashed lines represent the instantaneous temperature response based on enzyme kinetics.

The prediction that *b* should decline with temperature by 1.8% K^−1^ was consistent with the fitted regressions of the ratio of *R*_d,tg_ to *V*_cmax,tg_; we observed a small but significant negative response of *b* to growth temperature with a sensitivity of 2.0% ± 0.3%, while *b*_25_ was indeed independent of mGDD_0_, as predicted (Table 1, Fig. 2). The fitted temperature response of *R*_d,tg_ was consistently about 2% less steep than that of *V*_cmax,tg_ (Table 1).

The canonical value of *b*_25_ = 0.015, as assumed in the photosynthesis model of Collatz *et al.* (1991), was similar to the fitted value of *b*_25_ = 0.014 ± 0.001 based on the regression of *R*_d,25_ with respect to *V*_cmax,25_ (Table 2). The regression lines of *R*_d,tg_ with respect to *V*_cmax,tg_ when fitted to data in low (*T*_g_ < 15°C), medium (15°C < *T*_g_ < 25°C) and high (*T*_g_ > 25°C) temperature classes separately (Fig. S3) became shallower toward higher temperature classes, also consistent with the prediction of a negative response of *b* to temperature.

**Table 2:**
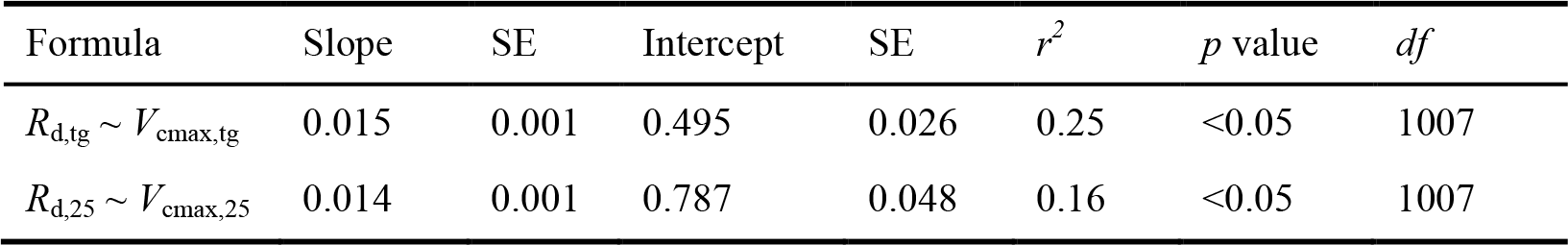
Summary statistics for Ordinary Least Squares regressions of area-based leaf dark respiration (*R*_d_) against area-based maximum carboxylation rate (*V*_cmax_) assessed at growth temperature and at 25°C. The fitted slopes and intercept are shown together with their standard errors (SE), the coefficient of determination (*r^2^*) and the degrees of freedom (*df*).

The difference between the enzyme-kinetic sensitivities of *R*_d_ and *V*_cmax_ to temperature implies that the sensitivity of acclimated *R*_d,tg_ to temperature (*β*_aR_) is 1.8% lower than that of *V*_cmax,tg_ (*β*_aV_), implying a theoretical optimum rate of increase of *R*_d,tg_ by 3.7% per degree. These theoretical responses are very close to those seen in the observations, but rather more shallow than the 9.9% and 8.1% K^−1^ predicted for the short-term responses of *V*_cmax_ and *R*_d_ from enzyme kinetics (Fig. 1). Theoretically predicted values of the fractional sensitivities of acclimated *R*_d,25_ (*β*_aR25_) and *V*_cmax,25_ (*β*_aV25_) to temperature are negative (−4.4 % K^−1^) and this is consistent with the observed negative responses of *R*_d,25_ and *V*_cmax,25_ to temperature seen in the data (Table 1). The observed negative response of *V*_cmax,25_ to growth temperature is identical (−4.4 ± 0.3% K^−1^) to our theoretical prediction; the observed response of 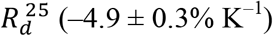 is marginally larger than the prediction.

Regressions performed without PPFD-normalization (Table S1) showed temperature responses with the same signs (positive for *V*_cmax_ and *R_d_* at growth temperature, negative at 25°C) but slightly steeper (positive slopes) or shallower (negative slopes) than in the main analyses – as expected due to the confounding of PPFD and temperature effects, which normalization removes. *R*^2^ values were consistently greater in the main analyses, by 6-7% for *V*_cmax_ and *R_d_* at growth temperature and 11-16% for *V*_cmax_ and *R_d_* at 25°C.

Thermal acclimation of *V*_cmax_ within PFTs was shown to be broadly consistent with the universal relationships evident in the whole dataset (Table S2). Consistent with Campbell et al. (2007) and Smith and Dukes (2017b), no significant differences in thermal acclimation were found between PFTs with different leaf phenology (evergreen versus deciduous), leaf form (needleleaf versus broadleaf), or life form (trees versus shrubs versus herbaceous plants), although the fitted slope for evergreen broadleaf trees is higher than predicted. Thermal acclimation of *R*_d_ within PFTs shows more variable results. Evergreen broadleaf trees show higher than predicted thermal sensitivities of both *R*_d_ and *V*_cmax_, but the fitted slope of their ratio versus growth temperature (−1.8% ± 0.4%) is identical with our predicted value (−1.8%).

Microclimatic acclimation (Niinemets & Keenan 2012), the likelihood that many measured leaves were at least partially shaded (Keenan & Niinemets 2017), and genetic variations involving different plant strategies may all have contributed to the within-site variations in *R*_d_ and *V*_cmax_ reflected in the vertical scatter of points in Fig. 2. The theory was applied here to predict variations in *R*_d_ and *V*_cmax_ across sites, however, and as much as 45% and 60% variation in the community-mean *R*_d,tg_ and *V*_cmax,tg_ respectively could indeed be explained by growth temperature – with responses in quantitative agreement with predictions (Table S3).

#### Relationships between dark respiration and carboxylation capacity

We examined the relationships between *R*_d_, *V*_cmax_ and other factors, in order to test our hypothesis that *R*_d_ is principally determined by *V*_cmax_. We found that measured *R*_d_ and *V*_cmax_ were positively correlated in the datasets when normalized either to mGDD_0_ (*R*^2^= 0.25) or to a reference temperature of 25°C (*R*^2^ = 0.16) (Table 2).

#### Relationships of dark respiration and photosynthetic capacity to other variables

Relationships of *R*_d_ and *V*_cmax_ to *N*_area_ were similar in strength when normalized to 25°C (*R*^2^ = 0.14 and 0.12) (Table 3), but notably weaker when considered at growth temperature (*R*^2^ = 0.05 for *R*_d,tg_ and 0.02 for *V*_cmax,tg_). LMA and *V*_cmax,25_ together accounted for 42% variation in *N*_area_, but most of this explanatory power coming from LMA (Table 3). Similarly, LMA and *R*_d,25_ together explained 41% variation in *N*_area_, but again most of this explanatory power is due to LMA (Table 3).

**Table 3:** Summary statistics for Ordinary Least Squares regressions of area-based leaf nitrogen content against area-based leaf dark respiration (*R*_d_), area-based leaf maximum capacity of carboxylation (*V*_cmax_) assessed at growth temprature and 25°C, and leaf mass per area (LMA). The fitted slopes (mean ± standard error) are shown together with the intercept (mean ± standard error), the adjusted coefficient of determination (*r^2^*) and the degrees of freedom (*df*). All variables were natural-log transformed.

| Coefficient | | | | | Intercept | $r^2$ | $df$ |
| --- | --- | --- | --- | --- | --- | --- | --- |
| $V_{\text{cmax,tg}}$ | $V_{\text{cmax},25}$ | $R_{d,tg}$ | $R_{d,25}$ | LMA | | | |
| 0.083±0.019 | | | | | 0.409±0.059 | 0.02 | 935 |
| 0.058±0.015 | | | | 0.491±0.021 | -1.849±0.107 | 0.39 | 934 |
| | 0.242±0.022 | | | | -0.199±0.081 | 0.12 | 935 |
| | 0.148±0.018 | | | 0.458±0.021 | -2.055±0.106 | 0.42 | 934 |
| | | 0.177±0.022 | | | 0.677±0.015 | 0.05 | 1165 |
| | | 0.107±0.018 | | 0.508±0.020 | -1.724±0.097 | 0.38 | 1164 |
| | | | 0.301±0.022 | | 0.590±0.013 | 0.14 | 1165 |
| | | | 0.174±0.019 | 0.471±0.020 | -1.606±0.097 | 0.41 | 1164 |

Considered on their own, neither LMA nor soil pH provided explanatory power in the variations of *R*_d_ and *V*_cmax_ (whether at 25°C and growth temperature). The inclusion of one or the other in addition to mGDD_0_ as a predictor provided negligible increases in explained variance (Table S4).

### Discussion

#### Acclimation of leaf dark respiration follows the acclimation of carboxylation capacity

Farquhar *et al.* (1980) modelled instantaneous *R*_d_ at 25°C as a fixed fraction (1.1%) of *V*_cmax_ at 25°C. Collatz et al. (1991) – citing Farquhar et al. (1980) – modelled *R*_d_ at 25°C as 1.5% of *V*_cmax_ at 25°C. Our results show that *R_d_* and *V*_cmax_ are indeed closely related, whether standardized to growth temperature or to 25°C, albeit with substantial scatter around the relationship. Moreover, the coefficient relating *R*_d,25_ to *V*_cmax,25 -_ estimated from this large global dataset (*b*_25_ = 0.014 ± 0.01, Table 2) is indistinguishable from the value of 0.015 used by Collatz *et al.* (1991). We also predict a slight negative response of *b* to temperature due to the difference in the kinetic responses of *R*_d_ and *V*_cmax_. This expectation is consistent with other studies (De Kauwe *et al.* 2016), and well supported by the finding here of a temperature effect in the regression of *b* (estimated from the *R*_d,tg_ to *V*_cmax,tg_ ratio), but not in that of *b*_25_ (estimated from the *R*_d,25_ to *V*_cmax,25_ ratio). The former provides a value of *β*_b_ close to our theoretical value of 1.8% K^−1^ (Table 1).

Atkin et al. (2015) indicated a decline in *b*_25_ with increasing growth temperature. Their dataset that did not included the LCE data. To compare our analysis with that of Atkin et al. (2015), we assessed the response of *b*_25_ to mGDD_0_ after excluding the LCE data. The response of *b*_25_ to mGDD_0_ became significant (*p* = 0.0076) but its sensitivity (*β_b25_* = −0.009 ± 0.003) was much lower than that of *b* (*β*_*b*_ = −0.024 ± 0.003, *p* < 0.001). Atkin *et al.* (2015) considered the mean temperature of the warmest three-month period as growth temperature (TWQ), whereas here all days with temperature above 0°C are accounted in the growing season – thus growth temperature by our definition (mGDD_0_) is lower than TWQ, except for the low and high ends of the scale (Fig. S4). This methodological difference might have contributed to a slightly different conclusion regarding the temperature sensitivity of *b*_25_.

Equation (1) would potentially allow predictions of the responses of *R*_d_ to other environmental determinants, including vapour pressure deficit, elevation and CO_2_, if these too are determined by the environmental responses of *V*_cmax_. However, the data currently available do not allow us to test these predictions, due to either the limited environmental range covered by the data (such as elevation or CO_2_) or the strong correlations between potential explanatory variables (including light and temperature, Fig. S2). Nevertheless, our theory provides a simple, first-principles approach to predicting the thermal acclimation of *R*_d_ – one of the most important mechanisms missing from current LSMs (Huntingford *et al.* 2017).

#### Observed temperature responses are consistent with optimal plant function

Many ecosystem models assume that *R*_d_ and *V*_cmax_ respond to temperature following the same functions that are routinely observed in short-term studies, disregarding acclimation. Our results contradict this assumption. Optimally acclimated values of both fluxes do increase with temperature, but much less steeply than expected from short-term responses. Instead it appears that leaves ‘discount’ enzyme-kinetic responses, so the two limiting photosynthetic rates remain similar *under the prevailing growth temperature*. Positive temperature responses arise because of the differential temperature sensitivities of two key quantities – the effective Michaelis-Menten coefficient of Rubisco (*K*) and the photorespiratory CO_2_ compensation point (*Γ*^*^) – in the Farquhar *et al.* (1980) model. These predictions are supported by the finding of temperature responses in the datasets evaluated here: that is, after the *R*_d_ and *V*_cmax_ values have been corrected to site-specific mean growing-season temperatures, we find responses closely similar to the predicted optimal responses (Fig. 2).

#### The acclimated sensitivity of dark respiration to warming

Acclimated *R*_d_ is predicted to increase with temperature by 3.7% per degree. This long-term (weekly or longer) sensitivity is supported by the data and is smaller than the enzyme-kinetic sensitivity of either *R*_d_ or *V*_cmax_. This response is generally conservative among PFTs.

Reich *et al.* (2016) conducted outdoor open-air warming experiments on ten boreal and temperate tree species. Whole plants were warmed by 3.4°C during four growing seasons, and showed even stronger acclimation of *R*_d_ and an acclimated sensitivity of only 1.5% per degree. Slot and Kitajima (2015) indicated a value of 5.7% for the temperature sensitivity of *R*_d_ based on a meta-analysis of 43 independent experiments (mostly short-term, laboratory studies). Our theory predicts *β*_aR_ as 3.7% per degree warming, intermediate between the values found by Reich *et al.* (2016) and Slot and Kitajima (2015). In a study that acclimated plants for seven days, Smith and Dukes (2017b) found even less acclimation of *R*_d_ than either of those studies. The differences among these three studies suggest that the time scale of exposure to elevated temperatures might play a role.

Heskel *et al.* (2016) presented data from different biomes that can be compared to our predictions of the acclimation of *R*_d_. We can estimate *a* in equation (3) by rearrangement, with *T*_ref_ = 25°C:

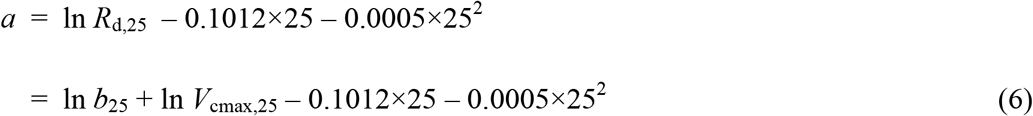

The thermal sensitivity of *a* should therefore be the same as that of *β*_aR25_. Its decline from tundra to tropical forests (Heskel *et al.*, 2016) is the result of *R*_d_ acclimation to growth temperature (Slot & Kitajima 2015; Vanderwel *et al.* 2015). Our theory estimates *a* independently: *a* = −2.502 when *V*_cmax,25_ = 50 µmol m^−2^ s^−1^ (i.e. equivalent, 25 = 1.4 µmol m^−2^ s^−1^), or *a* = −1.407 when *V*_cmax,25_ = 150 µmol m^−2^ s^−1^ (i.e. equivalent to *R*_d,25_ = 4.2 µmol m^−2^ s^−1^). These values are similar to those derived from observations by Heskel *et al.* (2016): lower for warm biomes (e.g. *a* = −2.749 for lowland tropical rainforest), where we would expect relatively low *V*_cmax,25_; and higher for cold biomes (e.g. *a* = −1.604 for tundra), where we would expect relatively high *V*_cmax,25_. These values from Heskel *et al.* (2016) allow us to approximate the thermal sensitivity of *a* as −4.6% K^−1^, assuming a growth temperature range of 25°C from tundra to rainforest − very close to our theoretical prediction of *β*_aR25_ = −4.4% K^−1^.

#### Relationships of respiration and photosynthetic capacity with leaf N

It is often assumed that *R*_d_ and *V*_cmax_ (assessed at standard temperature) are closely related to *N*_area_, and empirical studies have reported relationships of both quantities to *N*_area_ (Reich *et al.* 1998; Meir *et al.* 2001; Wright *et al.* 2004; Atkin *et al.* 2015). Correlation of both *R*_d,25_ and *V*_cmax,25_ with *N*_area_ is expected because of the significant fraction of *N*_area_ that is contained in Rubisco and other photosynthetic proteins, and the functional relationship between *R*_d_ and *V*_cmax_. (*N*_area_ should always be compared to *R*_d_ and *V*_cmax_ corrected to a common temperature, because the amount of “metabolic” N required to achieve a given catalytic activity is strongly dependent on temperature.) We found the expected positive relationships of *R*_d,25_ and *V*_cmax,25_ to *N*_area_, but they were not strong (*R*^2^ = 0.11 and 0.14, Table 3). Onoda *et al.* (2017) noted the substantial fraction of leaf N allocated to cell walls, in addition to the fraction contained in photosynthetic proteins. Our finding that far more variation in *N*_area_ can be explained by LMA than by *V*_cmax_ (Table 3) suggests that the structural (cell wall) component of *N*_area_ is important (see also (Dong *et al.* 2017). *N*_area_ is thus not a straightforward predictor of either *V*_cmax_ or *R*_d_, but rather contains both a metabolic component (related to *V*_cmax,25_) and a structural component proportional to LMA (Dong *et al.*, 2017). Mass-based quantities show a similar pattern (Table S5) with more variation in *N*_mass_ explained by LMA than by *V*_cmax_. However, LMA is negatively related to *N*_mass_, which indicates that the concentration of N in bulk leaf tissue is lower than that in the photosynthetic machinery.

The coordination hypothesis implies that *V*_cmax_, and therefore also the metabolic component of *N*_area_, should be determined by photosynthetic demand – rather than *V*_cmax_ being determined by *N*_area_, as is currently assumed in many models. This point is further addressed by (Maire *et al.* 2012; Dong *et al.* 2017), who noted that the co-ordination hypothesis predicts an optimal value for metabolic *N*_area_ on the basis of environmental conditions alone. The empirical *V*_cmax_−*N*_area_ relationship has been widely interpreted as a manifestation of ‘N limitation’ of photosynthesis at the leaf level (Luo *et al.* 2004), and underpins the use of *N*_area_ as a predictor of photosynthetic capacity in models (Ciais *et al.* 2014). An alternative interpretation is that leaf metabolic N depends primarily on the demand for photosynthetic capacity, which is set by the local environment and in turn determines the capacity for dark respiration; while N availability primarily influences the allocation of carbon to leaves versus other organs (LeBauer & Treseder 2008; Poorter *et al.* 2012).

### Conclusions

The observed thermal acclimation of *R*_d_ follows the optimization of *V*_cmax_ as predicted by the coordination hypothesis. This acclimation dampens the enzyme-kinetic response of *R*_d_ to temperature and shows little influence from other factors. The discrepancy between thermal acclimation and enzyme-kinetic thermal response implies that both *R*_d_ or *V*_cmax_, converted to 25°C or any other arbitrarily chosen reference temperature, must decline with plant growth temperature. These principles would be straightforward to incorporate in an LSM framework, as an alternative to PFT-based schemes in current use. The theory provides an explanation for the observed correlations between *N*_area_, *V*_cmax_ and *R*_d_ that differs from the common assumption that *N*_area_ determines *V*_cmax_ and *R*_d_, and supports an alternative perspective on the coupling between the terrestrial carbon and nitrogen cycles.

## References

Ainsworth, E.A. & Long, S.P. (2005). What have we learned from 15 years of free-air CO_2_ enrichment (FACE)? A meta-analytic review of the responses of photosynthesis, canopy properties and plant production to rising CO_2_. New Phytologist, 165, 351–372.

Amthor, J.S. (2000). The McCree–de Wit–Penning de Vries–Thornley respiration paradigms: 30 years later. Annals of botany, 86, 1–20.

Aspinwall, M.J., Drake, J.E., Campany, C., Vårhammar, A., Ghannoum, O., Tissue, D.T. et al. (2016). Convergent acclimation of leaf photosynthesis and respiration to prevailing ambient temperatures under current and warmer climates in Eucalyptus tereticornis. New Phytologist, 212, 354–367.

Atkin, O.K., Bloomfield, K.J., Bahar, N.H., Griffin, K.L., Heskel, M.A., Huntingford, C. et al. (2017). Leaf Respiration in Terrestrial Biosphere Models. In: Plant Respiration: Metabolic Fluxes and Carbon Balance (ed. Tcherkez, G). Springer-Nature: Netherlands, pp. 107–142.

Atkin, O.K., Bloomfield, K.J., Reich, P.B., Tjoelker, M.G., Asner, G.P., Bonal, D. et al. (2015). Global variability in leaf respiration in relation to climate, plant functional types and leaf traits. New Phytologist, 206, 614–636.

Atkin, O.K., Millar, A.H., Gardeström, P. & Day, D.A. (2000). Photosynthesis, carbohydrate metabolism and respiration in leaves of higher plants. In: Photosynthesis. Springer, pp. 153–175.

Atkin, O.K., Scheurwater, I. & Pons, T.L. (2007). Respiration as a percentage of daily photosynthesis in whole plants is homeostatic at moderate, but not high, growth temperatures. New Phytologist, 174, 367–380.

Atkin, O.K. & Tjoelker, M.G. (2003). Thermal acclimation and the dynamic response of plant respiration to temperature. Trends in plant science, 8, 343–351.

Bernacchi, C., Singsaas, E., Pimentel, C., Portis Jr, A. & Long, S. (2001). Improved temperature response functions for models of Rubisco - limited photosynthesis. Plant, Cell & Environment, 24, 253–259.

Blonder, B., Salinas, N., Bentley, L.P., Shenkin, A., Porroa, P.O.C., Tejeira, Y.V. et al. (2017). Predicting trait-environment relationships for venation networks along an Andes-Amazon elevation gradient. Ecology, 98, 1239–1255.

Booth, B.B., Jones, C.D., Collins, M., Totterdell, I.J., Cox, P.M., Sitch, S. et al. (2012). High sensitivity of future global warming to land carbon cycle processes. Environmental Research Letters, 7, 024002.

Bouma, T.J. (2005). Understanding plant respiration: separating respiratory components versus a process-based approach. In: Plant respiration. Springer, pp. 177–194.

Campbell, C., Atkinson, L., Zaragoza-Castells, J., Lundmark, M., Atkin, O. & Hurry, V. (2007). Acclimation of photosynthesis and respiration is asynchronous in response to changes in temperature regardless of plant functional group. New Phytologist, 176, 375–389.

Cannell, M. & Thornley, J. (2000). Modelling the components of plant respiration: some guiding principles. Annals of Botany, 85, 45–54.

Ciais, P., Sabine, C., Bala, G., Bopp, L., Brovkin, V., Canadell, J. et al. (2014). Carbon and other biogeochemical cycles. Climate change 2013: the physical science basis. Contribution of Working Group I to the Fifth Assessment Report of the Intergovernmental Panel on Climate Change, 465–570.

Collatz, G.J., Ball, J.T., Grivet, C. & Berry, J.A. (1991). Physiological and environmental regulation of stomatal conductance, photosynthesis and transpiration: a model that includes a laminar boundary layer. Agricultural and Forest meteorology, 54, 107–136.

Collatz, G.J., Berry, J.A., Farquhar, G.D. & Pierce, J. (1990). The relationship between the Rubisco reaction mechanism and models of photosynthesis. Plant, Cell & Environment, 13, 219–225.

Cowan, I.R. (1986). Economics of carbon fixation in higher plants. In: On the economy of plant form and function (ed. Givnish, TJ). Cambridge University Press Cambridge, pp. 133–170.

Cox, P.M., Betts, R.A., Jones, C.D., Spall, S.A. & Totterdell, I.J. (2000). Acceleration of global warming due to carbon-cycle feedbacks in a coupled climate model. Nature, 408, 184–187.

Davis, T.W., Prentice, I.C., Stocker, B.D., Thomas, R.T., Whitley, R.J., Wang, H. et al. (2017). Simple process-led algorithms for simulating habitats (SPLASH v. 1.0): robust indices of radiation, evapotranspiration and plant-available moisture. Geoscientific Model Development, 10, 689.

De Kauwe, M.G., Lin, Y.-S., Wright, I.J., Medlyn, B.E., Crous, K.Y., Ellsworth, D.S. et al. (2016). A test of the ‘one-point method’ for estimating maximum carboxylation capacity from field-measured, light-saturated photosynthesis. New Phytologist, 210, 1130–1144.

Dewar, R., Mauranen, A., Mäkelä, A., Hölttä, T., Medlyn, B. & Vesala, T. (2018). New insights into the covariation of stomatal, mesophyll and hydraulic conductances from optimization models incorporating nonstomatal limitations to photosynthesis. New Phytologist, 217, 571–585.

Dong, N., Prentice, I.C., Evans, B.J., Caddy-Retalic, S., Lowe, A.J. & Wright, I.J. (2017). Leaf nitrogen from first principles: field evidence for adaptive variation with climate. Biogeosciences, 14, 481–495.

Drake, J.E., Tjoelker, M.G., Aspinwall, M.J., Reich, P.B., Barton, C.V., Medlyn, B.E. et al. (2016). Does physiological acclimation to climate warming stabilize the ratio of canopy respiration to photosynthesis? New Phytologist, 211, 850–863.

Farquhar, G.D., Buckley, T.N. & Miller, J.M. (2002). Optimal stomatal control in relation to leaf area and nitrogen content. Silva Fennica, 36, 625–637.

Farquhar, G.D., von Caemmerer, S.v. & Berry, J. (1980). A biochemical model of photosynthetic CO_2_ assimilation in leaves of C_3_ species. Planta, 149, 78–90.

Friedlingstein, P., Meinshausen, M., Arora, V.K., Jones, C.D., Anav, A., Liddicoat, S.K. et al. (2014). Uncertainties in CMIP5 climate projections due to carbon cycle feedbacks. Journal of Climate, 27, 511–526.

Gifford, R.M. (2003). Plant respiration in productivity models: conceptualisation, representation and issues for global terrestrial carbon-cycle research. Functional Plant Biology, 30, 171–186.

Givnish, T.J. (1986a). On the economy of plant form and function. Cambridge University Press, Cambridge.

Givnish, T.J. (1986b). Optimal stomatal conductance, allocation of energy between leaves and roots, and the marginal cost of transpiration. In: On the economy of plant form and function (ed. Givnish, TJ). Cambridge University Press Cambridge, pp. 25–55.

Gobron, N., Pinty, B., Taberner, M., Mélin, F., Verstraete, M. & Widlowski, J.-L. (2006). Monitoring the photosynthetic activity of vegetation from remote sensing data. Advances in Space Research, 38, 2196–2202.

Harrison, S.P., Prentice, I.C., Barboni, D., Kohfeld, K.E., Ni, J. & Sutra, J.P. (2010). Ecophysiological and bioclimatic foundations for a global plant functional classification. Journal of Vegetation Science, 21, 300–317.

Haxeltine, A. & Prentice, I.C. (1996). A general model for the light-use efficiency of primary production. Functional Ecology, 10, 551–561.

Heskel, M.A., O’Sullivan, O.S., Reich, P.B., Tjoelker, M.G., Weerasinghe, L.K., Penillard, A. et al. (2016). Convergence in the temperature response of leaf respiration across biomes and plant functional types. Proceedings of the National Academy of Sciences, 113, 3832–3837.

Hoefnagel, M., Atkin, O. & Wiskich, J. (1998). Interdependence between chloroplasts and mitochondria in the light and the dark. Biochimica Et Biophysica Acta-Bioenergetics, 1366, 235–255.

Huntingford, C., Atkin, O.K., Heskel, M.A., Martinez-de la Torre, A., Harper, A.B., Bloomfield, K.J. et al. (2017). Implications of improved representation of plant respiration in a changing climate. Nature Communications, 8, 1602.

Huntingford, C., Zelazowski, P., Galbraith, D., Mercado, L.M., Sitch, S., Fisher, R. et al. (2013). Simulated resilience of tropical rainforests to CO_2_-induced climate change. Nature Geoscience, 6, 268.

Kattge, J. & Knorr, W. (2007). Temperature acclimation in a biochemical model of photosynthesis: a reanalysis of data from 36 species. Plant, cell & environment, 30, 1176–1190.

Keenan, T.F. & Niinemets, Ü. (2017). Global leaf trait estimates biased due to plasticity in the shade. Nature plants, 3, 16201.

Keenan, T.F., Prentice, I.C., Canadell, J.G., Williams, C.A., Wang, H., Raupach, M. et al. (2016). Recent pause in the growth rate of atmospheric CO_2_ due to enhanced terrestrial carbon uptake. Nature communications, 7, 13428.

Kelley, D., Prentice, I.C., Harrison, S., Wang, H., Simard, M., Fisher, J. et al. (2013). A comprehensive benchmarking system for evaluating global vegetation models. Biogeosciences, 10, 3313–3340.

Kikuzawa, K., Onoda, Y., Wright, I.J. & Reich, P.B. (2013). Mechanisms underlying global temperature-related patterns in leaf longevity. Global Ecology and Biogeography, 22, 982–993.

Lin, Y.-S., Medlyn, B.E., Duursma, R.A., Prentice, I.C., Wang, H., Baig, S. et al. (2015). Optimal stomatal behaviour around the world. Nature Climate Change, 5, 459–464.

Luo, Y., Su, B., Currie, W.S., Dukes, J.S., Finzi, A., Hartwig, U. et al. (2004). Progressive nitrogen limitation of ecosystem responses to rising atmospheric carbon dioxide. AIBS Bulletin, 54, 731–739.

Maire, V., Martre, P., Kattge, J., Gastal, F., Esser, G., Fontaine, S. et al. (2012). The coordination of leaf photosynthesis links C and N fluxes in C_3_ plant species. PloS one, 7, e38345.

Marquet, P.A., Allen, A.P., Brown, J.H., Dunne, J.A., Enquist, B.J., Gillooly, J.F. et al. (2014). On theory in ecology. BioScience, 64, 701–710.

Meir, P., Grace, J. & Miranda, A. (2001). Leaf respiration in two tropical rainforests: constraints on physiology by phosphorus, nitrogen and temperature. Functional Ecology, 15, 378–387.

New, M., Hulme, M. & Jones, P. (2000). Representing Twentieth-Century Space-Time Climate Variability. Part II: Development of 1901-96 Monthly Grids of Terrestrial Surface Climate. Journal of Climate, 13, 2217–2238.

Niinemets, Ü. & Keenan, T. (2012). Measures of light in studies on light-driven plant plasticity in artificial environments. Frontiers in plant science, 3, 156.

Noguchi, K. & Yoshida, K. (2008). Interaction between photosynthesis and respiration in illuminated leaves. Mitochondrion 8, 87–99.

Onoda, Y., Wright, I.J., Evans, J.R., Hikosaka, K., Kitajima, K., Niinemets, Ü. et al. (2017). Physiological and structural tradeoffs underlying the leaf economics spectrum. New Phytologist, 214, 1447–1463.

Reich, P.B., Sendall, K.M., Stefanski, A., Wei, X., Rich, R.L. & Montgomery, R.A. (2016). Boreal and temperate trees show strong acclimation of respiration to warming. Nature, 531, 633–636.

Reich, P.B., Walters, M.B., Ellsworth, D.S., Vose, J.M., Volin, J.C., Gresham, C. et al. (1998). Relationships of leaf dark respiration to leaf nitrogen, specific leaf area and leaf life-span: a test across biomes and functional groups. Oecologia, 114, 471–482.

Rogers, A. (2014). The use and misuse of V_c,max_ in Earth System Models. Photosynthesis Research, 119, 15–29.

Scafaro, A.P., Xiang, S., Long, B.M., Bahar, N.H., Weerasinghe, L.K., Creek, D. et al. (2017). Strong thermal acclimation of photosynthesis in tropical and temperate wet-forest tree species: the importance of altered Rubisco content. Global Change Biology, 23, 2783–2800.

Slot, M. & Kitajima, K. (2015). General patterns of acclimation of leaf respiration to elevated temperatures across biomes and plant types. Oecologia, 177, 885–900.

Smith, N.G. & Dukes, J.S. (2013). Plant respiration and photosynthesis in global-scale models: incorporating acclimation to temperature and CO_2_. Global Change Biology, 19, 45–63.

Smith, N.G. & Dukes, J.S. (2017a). LCE: leaf carbon exchange data set for tropical, temperate, and boreal species of North and Central America. Ecology, 98, 2978–2978.

Smith, N.G. & Dukes, J.S. (2017b). Short-term acclimation to warmer temperatures accelerates leaf carbon exchange processes across plant types. Global change biology, 23, 4840–4853.

Smith, N.G., Malyshev, S.L., Shevliakova, E., Kattge, J. & Dukes, J.S. (2016). Foliar temperature acclimation reduces simulated carbon sensitivity to climate. Nature Climate Change, 6, 407–411.

Tcherkez, G., Boex-Fontvieille, E, Mahe, A, Hodges, M (2012). Respiratory carbon fluxes in leaves. Current Opinion in Plant Biology 15, 308–314.

Tilman, D. (1999). The ecological consequences of changes in biodiversity: a search for general principles. Ecology, 80, 1455–1474.

Togashi, F.H., Prentice, I.C., Atkin, O.K., Macfarlane, C., Prober, S.M., Bloomfield, K.J. et al. (2018). Thermal acclimation of leaf photosynthetic traits in an evergreen woodland, consistent with the co-ordination hypothesis. Biogeosciences, 15, 3461–3474.

Vanderwel, M.C., Slot, M., Lichstein, J.W., Reich, P.B., Kattge, J., Atkin, O.K. et al. (2015). Global convergence in leaf respiration from estimates of thermal acclimation across time and space. New Phytologist, 207, 1026–1037.

Wang, H., Prentice, I. & Davis, T. (2014). Biophsyical constraints on gross primary production by the terrestrial biosphere. Biogeosciences, 11, 5987–6001.

Wang, H., Prentice, I.C., Davis, T.W., Keenan, T.F., Wright, I.J. & Peng, C. (2017a). Photosynthetic responses to altitude: an explanation based on optimality principles. New Phytologist, 213, 976–982.

Wang, H., Prentice, I.C., Keenan, T.F., Davis, T.W., Wright, I.J., Cornwell, W.K. et al. (2017b). Towards a universal model for carbon dioxide uptake by plants. Nature Plants, 3, 734–741.

Weng, E., Farrior, C.E., Dybzinski, R. & Pacala, S.W. (2017). Predicting vegetation type through physiological and environmental interactions with leaf traits: Evergreen and deciduous forests in an earth system modeling framework. Global change biology, 23, 2482–2498.

Wolf, A., Anderegg, W.R. & Pacala, S.W. (2016). Optimal stomatal behavior with competition for water and risk of hydraulic impairment. Proceedings of the National Academy of Sciences, 113, E7222–E7230.

Wright, I.J., Reich, P.B., Atkin, O.K., Lusk, C.H., Tjoelker, M.G. & Westoby, M. (2006). Irradiance, temperature and rainfall influence leaf dark respiration in woody plants: evidence from comparisons across 20 sites. New Phytologist, 169, 309–319.

Wright, I.J., Reich, P.B. & Westoby, M. (2003). Least-cost input mixtures of water and nitrogen for photosynthesis. The American Naturalist, 161, 98–111.

Wright, I.J., Reich, P.B., Westoby, M., Ackerly, D.D., Baruch, Z., Bongers, F. et al. (2004). The worldwide leaf economics spectrum. Nature, 428, 821–827.

Xu, X., Medvigy, D., Joseph Wright, S., Kitajima, K., Wu, J., Albert, L.P. et al. (2017). Variations of leaf longevity in tropical moist forests predicted by a trait-driven carbon optimality model. Ecology letters, 20, 1097–1106.

Ziehn, T., Kattge, J., Knorr, W. & Scholze, M. (2011). Improving the predictability of global CO_2_ assimilation rates under climate change. Geophysical Research Letters, 38, L10404.

